# Potential intrahospital dissemination of *Pseudomonas aeruginosa* carrying the *bla*_IMP-1_ gene within a Tn*7*-like transposon

**DOI:** 10.1101/2023.10.06.561298

**Authors:** Lin Zheng, Zixian Wang, Jingyi Guo, Jiayao Guan, Quanliang Li, Gejin Lu, Jie Jing, Shiwen Sun, Yang Sun, Xue Ji, Bowen Jiang, Ping Chen, Yanling Yang, Lingwei Zhu, Xuejun Guo

**Affiliations:** Key Laboratory of Jilin Province for Zoonosis Prevention and Control, Changchun Veterinary Research Institute, Chinese Academy of Agriculture Sciences, Changchun, Jilin 130122, China; Institute of Special Animal and Plant Science of Chinese Academy of Agricultural Science, Changchun, Jilin 130112, China; College of Veterinary Medicine, Jilin Agriculture University, Changchun, Jilin 130118, China; School of Food and Engineering, Jilin Agriculture University, Changchun, Jilin 130118 China; The second Norman Bethune hospital, Jilin University, Changchun, Jilin 130041, China

**Author notes:** Address correspondence to Xuejun Guo, Lingwei Zhu, or Yanling Yang,. Lin Zheng and Zixian Wang contributed equally to this work. Author order was determined by the corresponding author after negotiation.

**Keywords:** *Pseudomonas aeruginosa*, *bla*_IMP-1_, antimicrobial resistance, transposon

## Abstract

We aimed to determine the molecular characteristics of carbapenem-resistant *Pseudomonas aeruginosa* 18081308 and 18083286 isolated from the urine and sputum of two Chinese patients respectively, and analyzed the formation mechanism of the genetic environment in which it carries *bla*_IMP-1_. Bacterial genome sequencing was carried out on strains 18081308 and 18083286 to obtain their whole genome sequence. Average nucleotide identity (ANI) was used for their precise species identification. Serotyping and multilocus sequence typing were performed. Furthermore, the acquired resistance genes, and virulence factors of these strains were identified. The carbapenem-resistant *P. aeruginosa* strains isolated in the present study were of sequence type ST865 and serotype O6. They all carried the same virulence factors (PLC, ExoSTY) and resistance genes (*aacC2*, *tmrB*, and *bla*_IMP-1_). Tn*6411*, a Tn*7*-like transposon carrying *bla*_IMP-1_, was found in both strains. Detailed genetic dissection was applied to this transposon to display the genetic environment of *bla*_IMP-1_. The *aacC2*-*tmrB* region remnant-Tn*6411* backbone was the original structure of this type of transposon. A Tn*402*-like type 1 integron (*intl1*-*aac(6’)-II*-*bla*_IMP-1_) was inserted into it and formed a stable structure, which was localized in the chromosome by TnsD for transmission within *P. aeruginosa*; the original structure of Tn*7*-like transposon was localized on the plasmid by TnsE for horizontal transmission between bacterial species.

The intrahospital dissemination of *P. aeruginosa* ST865 isolated in this study was episodic. The *bla*_IMP-1_-carrying Tn*7*-like transposon might enhance their ability to survive under drug selection pressure and aggravate the difficulty in treating infections.

## Introduction

*Pseudomonas aeruginosa* is a zoonotic opportunistic bacterial pathogen that is ubiquitous in diverse environments, including water, animal-related food, the surface of medical instruments, and sewage systems in hospitals ^1,2^. It carries a large variety of virulence factors and can cause bacteremia, ventilator-associated pneumonia, cystic fibrosis, and chronic obstructive pulmonary disease. It has the ability to form biofilms and attach to the surface of medical instruments and food ^1^. It can spread in healthcare settings from one person to another through contaminated hands or surfaces. It is easily disseminated within hospitals. It caused an estimated 32,600 infections among hospitalized patients and 2,700 estimated deaths in the United States according to the Threat Estimate 2019 report ^3^. Clinically, *P. aeruginosa* infection is usually treated by antimicrobial therapy, and prolonged use of antibiotics to achieve bacterial cure is commonly practiced ^4^. Resistance genes can be acquired by the transfer of mobile genetic elements (such as plasmids, transposons, and integrative and conjugative elements) among bacterial strains, resulting in the development of multidrug-resistant *P. aeruginosa* in chronically infected patients ^5,6^. Carbapenems are the most important antibiotics for treating multidrug-resistant bacterial infections, but *P. aeruginosa* is currently also resistant to carbapenems due to the acquisition of carbapenemases, among other reasons, impeding treatment.

In the present study, two *P. aeruginosa* isolates (18081308 and 18083286) were obtained from two patients (urine and sputum samples, respectively) admitted to a public hospital (Changchun, China) in August 2018. Their whole genome sequences and molecular characteristics were determined. Both isolates belonged to the same multilocus sequence type (ST865) and serotype (O6). They both carried *bla*_IMP-1_. Detailed genetic dissection was applied to a Tn*7*-like transposon carrying *bla*_IMP-1_ to display its genetic environment. The data presented here provide a deeper understanding of drug resistance gene acquisition in *P. aeruginosa* from a genomic and bioinformatic point of view.

## Materials and methods

### Bacterial isolation and identification

Strains 18081308 and 18083286 were isolated from urine and sputum specimens, respectively, from two patients in a public hospital in China in 2018. The species was determined based on the partial sequence of the 16S rRNA gene ^7^.

Minimum inhibitory concentrations (MICs) of amikacin, gentamicin, meropenem, imipenem, cefazolin, ceftazidime, cefotaxime, cefepime, aztreonam, ampicillin, piperacillin, amoxicillin-clavulanate, ampicillin-sulbactam, piperacillin-tazobactam, trimethoprim-sulfamethoxazole, chloramphenicol, ciprofloxacin, levofloxacin, moxifloxacin, and tetracycline were tested by BD Phoenix-100, using *Escherichia coli* ATCC25922 as a control. Drug resistance and sensitivity were judged based on the Clinical and Laboratory Standards Institute guidelines.

### Sequencing and sequence assembly

Bacterial genomic DNA was extracted from strains 18081308 and 18083286 using the UltraClean Microbial Kit and sequenced using an Illumina NovaSeq PE150 platform. Trimmomatic V10 was used to remove the PCR adapters and low-quality reads, and SPAdes (http://cab.spbu.ru/software/spades/) was used for sequence assembly ^8^. Strain 18083286 was randomly selected for another DNA extraction step, and the newly obtained DNA was sequenced by a PacBio RSII sequencer.

### Sequence annotation and comparison

Precise species identification was performed by pairwise average nucleotide id-entity (ANI) (http://www.ezbiocloud.net/tools/ani) analysis between genome sequenc-es and the *P. aeruginosa* reference genome PAO1 (GenBank ID: NC_002516.2). An ≥95% ANI cut-off was used to define bacterial species ^9^. PAst (https://cge.foo-d.dtu.dk/services/PAst/) was used to perform serotyping. RAST *2.0* ^10^ and BLASTP /BLASTN ^11^searches were conducted to predict open reading frames (ORFs), onlin-e databases CARD ^12^(https://card.mcmaster.ca/), ResFinder 4.0 ^13^(https://cge.cbs.dtu. dk/services/ResFinder/), and VFDB ^14^(http://www.mgc.ac.cn/VFs/) were used to iden-tify resistance genes and virulence genes, and ISfinder ^15^(https://www-is.biotoul.fr/; Lastest Database Update 2021-9-21), TnCentral (https://tncentral.ncc.unesp.br), INT EGRALL (http://integrall.bio.ua.pt/) ^16^, and ICEberg 2.0 (http://db-mml.sjtu.edu.cn/IC Eberg/) ^17^were used to identify mobile elements. Pairwise sequence comparisons w-ere carried out by BLASTN. Gene organization diagrams were drawn by Inkscape 1.0 (http://inkscape.org/en/).

Multilocus sequence types (STs) were obtained by uploading their genomes (including their seven conserved housekeeping genes *acsA*, *aroE*, *gtaA*, *mutL*, *nuoD*, *ppsA*, and *trpE*) to pubMLST (https://pubmlst.org/).

### Conjugation experiments

Conjugation experiments were performed as described previously ^18^. Briefly, strains 18083286 and 18081308 were used as donors, and the sodium azide-resistant *E. coli* J53 was used as a recipient. Donor and recipient strains were separately cultured overnight at 37 ℃. Then, 3 mL of 18083286 culture was mixed with an equal volume of *E. coli* J53 culture. The mixed cells were harvested by centrifugation for 3 min at 12,000 ×g, washed with 3 mL of lysogeny broth (LB), and resuspended in 150 µL of LB. The mixture was spotted on a 1-cm^2^ hydrophilic nylon membrane filter (Millipore) with a 0.45-µm pore size, which was then placed on an LB agar plate and incubated for mating at 37 ℃ for 6 h. The cells were recovered from the filter membrane and spotted on LB agar containing 100 µg/mL sodium azide and 4 µg/mL imipenem to select carbapenem-resistant *E. coli* transconjugants.

### Nucleotide sequence accession numbers

The contig sequences of strains 18081308 and 18083286 have been submitted to GenBank under accession numbers GCA_024718375.1 and GCA_024714245.1. The complete sequence of 18083286 has been submitted to GenBank under accession number CP110368.

### Results and discussion

After strains 18083286 and 18081308 were cultured overnight at 37 ℃, round flat colonies with non-regular edges, fusion growth, pyocyanin production, and the presence of metallic sheen were found on brain heart infusion agar (imipenem concentration: 4 µg/mL).

Strains 18083286 and 18081308 were identified as *P. aeruginosa* by the BD Phoenix-100 identification system and based on the 16S rRNA gene. Table 1 shows the drug resistance spectrum of strain 18083286. After Illumina NovaSeq PE150 sequencing (basic information about the Illumina sequencing results is provided in Table 2), it was found that their ANI values were more than 95% with the reference strain *P. aeruginosa* PAO1 (GenBank ID: NC_002516.2), and they were confirmed to be *P. aeruginosa* (ANI values of *P. aeruginosa* 18083286 and 18081308 are provided in Table S1).

**TABLE 1.** Antimicrobial susceptibility of strain 18083286.

| Antimicrobial type | Antimicrobial | MIC (μg/mL)<sup>a</sup> | SIR<sup>b</sup> |
| --- | --- | --- | --- |
| Aminoglycoside | Amikacin | 16 | S |
|  | Gentamicin | >8 | R |
| β-lactam | Imipenem | 8 | R |
|  | Meropenem | 8 | R |
|  | Cefazolin | >16 | R |
|  | Ceftazidime | >16 | R |
|  | Cefotaxime | >32 | R |
|  | Cefepime | >16 | R |
|  | Aztreonam | 8 | S |
|  | Ampicillin | >16 | R |
|  | Piperacillin | 16 | S |
|  | Amoxicillin-clavulanate | >16/8 | R |
|  | Ampicillin-sulbactam | >16/8 | R |
|  | Piperacillin-tazobactam | 32/4 | I |
| Colistin | Colistin | 1 | NA |
| Sulfonamide | Trimethoprim-sulfamethoxazole | 2/38 | R |
| Chloramphenicol | Chloramphenicol | >16 | R |
| Quinolones | Ciprofloxacin | ≤0.5 | S |
|  | Levofloxacin | ≤1 | S |
|  | Moxifloxacin | 4 | NA |
| Tetracycline | Tetracycline | 8 | R |

<sup>a</sup>MIC, minimum inhibitory concentration.
<sup>b</sup>SIR, Susceptible (S), intermediate (I), resistant (R).
NA, not applicable.

**TABLE 2.**
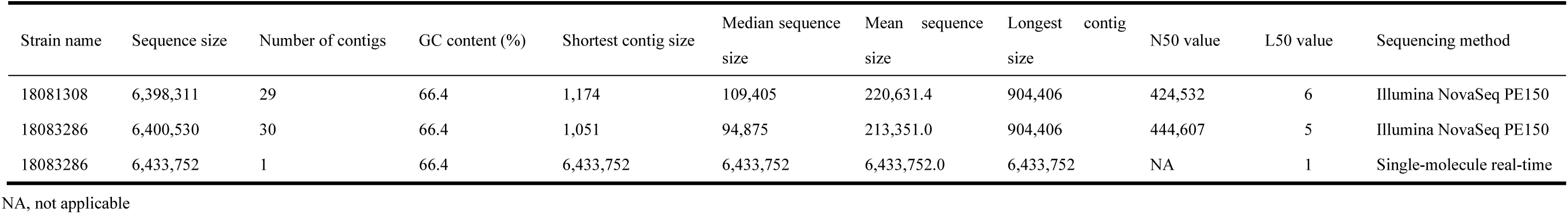
Basic information about bacterial sequencing results.

Both isolates belonged to the same multilocus sequence type (ST865) and serotype (O6) based on MLST and PAst screening. There are 20 serotypes of *P. aeruginosa*, of which serotype O6 is one of the most common ^19^. ST865 is not a pandemic clonal group. Until 2022, the MLST database contained a total of four strains of *P. aeruginosa* ST865: strains 18081308 and 18083286 isolated in this study, strain AZPAE14882 of unknown origin, and strain AUS151 isolated from soft tissue in Australia in 2008. Therefore, ST865 transmission in the hospital may only be an incidental event due to incomplete hospital disinfection.

It was found that both strains carry genes encoding virulence factors phospholipase C (PLC) and exotoxins (ExoS, ExoT, and ExoY). PLC is a thermolabile hemolysin. It can degrade the phospholipid surfactant, which functions to reduce the surface tension so that the alveoli do not collapse completely when the air leaves them during breathing ^20,21^. ExoS, ExoT, and ExoY are secreted by the type III secretion system (T3SS). They can disrupt the cytoskeleton, induce host cell rounding, disrupt intercellular tight junctions, prevent wound healing, and inhibit bacterial internalization into epithelial cells and macrophages ^22,23^. Genes for the most pathogenic virulence factors ExoU (a T3SS effector) and ExoA ^24–26^, were not found in this study. Consistent with a previous report ^27^, the strains did not carry both ExoS and ExoU, so strains in this study might cause cellular damage in immunocompromised patients, but do not cause acute toxicity.

They were resistant to many antibiotics in addition to imipenem, including: gentamicin, meropenem, cefazolin, ceftazidime, cefotaxime, cefepime, ampicillin, amoxicillin-clavulanate, ampicillin-sulbactam, trimethoprim-sulfamethoxazole, chloramphenicol, and tetracycline. The results are summarized in Table 1. Aminoglycoside resistance genes (*aac(6’’)-II*, *aac(3)-IId*, and *aph(3’’)-IIb*), an amphenicol resistance gene (*catB7*), β-lactam resistance genes (*bla*_OXA-486_, *bla*_IMP-1_, and *bla*_PAO_), and a fosfomycin resistance gene (*fosA*) were identified by ResFinder in these strains.

Since the two strains had the same resistance profile, ST type, serotype, acquired resistance genes, and virulence factors, and their ANI value was 99.99% ^28^(ANI values are provided in Table S1), one of the two strains was randomly selected for genetic environment analysis of the carbapenem resistance gene *bla*_IMP-1_. Single-molecule real-time (SMRT) sequencing (basic information about SMRT sequencing results is provided in Table 2) showed that the chromosome of strain 18083286 was 6.4 Mb and its GC content was 66.3%; no plasmid was detected. *bla*_IMP-1_ was located in a Tn*7*-like transposon in the bacterial chromosome.

A 37.53-kb transposon was inserted at a *glmS* (glucosamine-fructose-6-phosphate aminotransferase) site in the chromosome of *P. aeruginosa* 18083286. It had a complete set of Tn*7*-family core transposon-encoded proteins (TnsABCDE), but with very low levels of nucleotide identity with Tn*7* counterparts. This structure had the closest phylogenetic relationship with Tn*6411* (a Tn*7*-like family transposon) in *P. aeruginosa* 12939 (GenBank ID: CP024477.1; coverage: 100%, identity: 100%); thus, Tn*7*-like transposon was identified as Tn*6411*. It was first discovered in a *P. aeruginosa* strain from China in 2018 ^29^.

Until October 2022, only 10 transposons with the same TnsA as Tn*6411* were indexed in GenBank (Table 3 shows strain information). Among them, strains 12939, 18083286, and HB2011305RE, and plasmid p201330 carried Tn*6411*, and others carried its derived structures, named Tn6*411*-like. They mainly included *P. aeruginosa* from China, but *Acinetobacter spp.* from Sydney, Australia, and *P. aeruginosa* from India were also found. Except for Tn*6411*_18083286_, Tn*6411*-like_SE5430_, and Tn*6411*-like_PA34_ transposons, others contained 20 bp TnsB-binding sites plus 26 bp inverted repeats (IRs), which were terminal flanking regions (Table S2). They contained complete TnsABC+TnsD/E proteins, which were encoded by genes inserted into attTn*7* or plasmids capable of transfer between bacteria. Except for Tn*6411*-like_PA34_ (Indian), all others carried a truncated *aacC2*-*tmrB* region. The intact structure (IS*26-aacC2-tmrB* region remnant*-bla*_TEM-1_) was found in pEl15573 ^30,31^. It was derived from transposon Tn*2.* The structure (IS*26-aacC2-tmrB* region remnant) was found in the IncR/IncP6 fusion plasmid pCRE3-KPC carried by *Citrobacter braakii* ^32^. The truncated *aacC2*-*tmrB* region identified in this study was likely to be an intact structure from which first *bla*_TEM-1_ and then IS*26* was deleted.

**TABLE 3.**
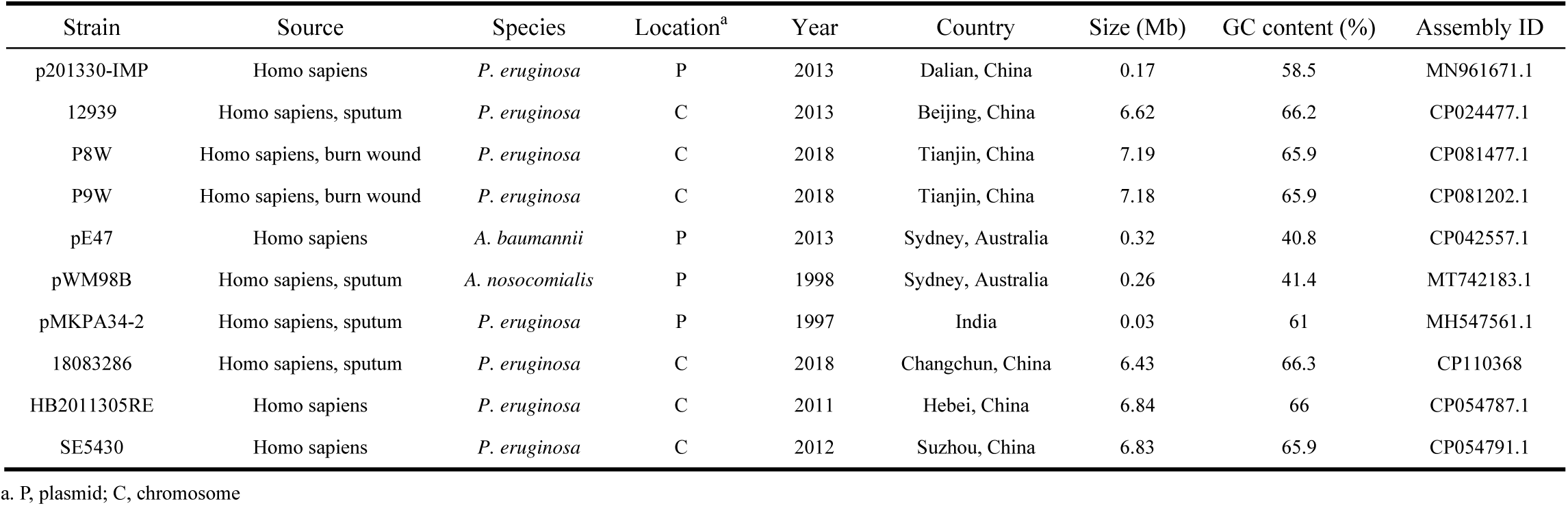
The information of strains carrying Tn*6411* and its derived structures.

All ten Tn*6411* transposons (until February 2022) listed in GenBank are shown in Figure 1, an integron carrying *bla*_IMP-1_ (a carbapenem resistance gene) and *aac(6’’)-II* (an aminoglycoside resistance gene) named In992 was inserted between *pinR* (encoding a DNA site-specific recombinase) and a gene (encoding a methyltransferase domain protein) that serves the backbone of Tn*6411*_12939_ (Beijing, China), Tn*6411*_18083286_ (Changchun, China), Tn*6411*_HB2011305RE_ (Changchun, China), and Tn*6411*_p201330_ (Changchun, China). One DR copy (ATGCCCGC) of In992 was found upstream of *pinR* (Tn*6411*-like_P8W_, Tn*6411*-like_P9W_, and Tn*6411*-like_SE5430_), which enabled the insertion of In992, resulting in a bilateral DR sequence (ATGCCCGC). Tn*6411* was carried by other strains that do not carry In992 or other integrons. The truncated Tn*402* transposition module in In992 had undergone the deletion of the partial TniA sequence and the complete TniBQR sequence and lost its self-transfer capability, making it stable in Tn*6411 and* Tn*6411*-like transposon.

**Figure 1.**
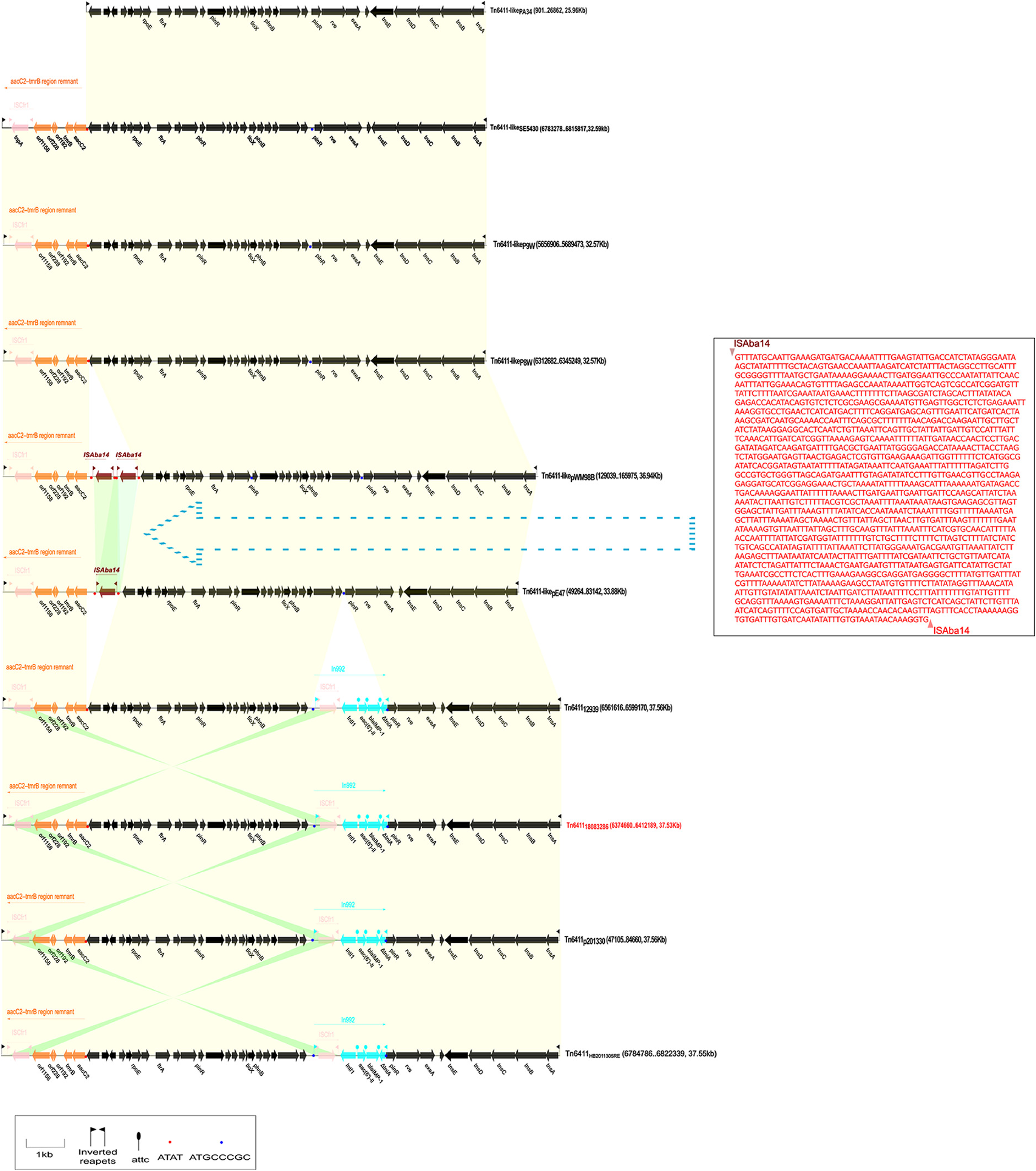
Linear alignment map of Tn*6411* and its derived structures. The backbone region is shown in black, In992 is shown in light blue, the *aacC2*-*tmrB* region remnant is shown in orange, IS*Aba14* is shown in brownish red, and IS*Cfr1* is shown in pink. The shaded region represents a region with >90% nucleotide identity. All transposons encoded complete TnsABC+TnsD/E proteins. Except for Tn*6411*-like_PA34_, all others carried a truncated *aacC2*-*tmrB* region remnant. An integron named In992 carrying *bla*_IMP-1_ and *aac(6’’)-II* was inserted between *pinR* and a gene encoding a methyltransferase domain protein from the backbone of Tn*6411*_12939_, Tn*6411*_18083286_, Tn*6411*_HB2011305RE_, Tn*6411*_p201330_, and Tn*6411* carried by other strains that do not carry In992 or other integrons. The truncated Tn*402* transposition module in In992 had undergone the deletion of the partial TniA sequence. Although Tn*6411*-like_pE47_ and Tn*6411*-like_pWM98B_ from *A. baumannii* and *A. nosocomialis* did not carry In992, one or two copies of IS*Aba14* were inserted downstream of the *aacC2*-*tmrB* region remnant. Tn*6411*-like_pE47_ had a one-copy IS*Aba14* difference from Tn*6411*-like_pWM98B_ in addition to a 1777-bp sequenSce difference.

One or two copies of IS*Aba14* were inserted upstream of the *aacC2*-*tmrB* region remnant, although Tn*6411*-like_pE47_ and Tn*6411*-like_pWM98B_ from *Acinetobacter baumannii* (Sydney, Australia) and *Acinetobacter nosocomialis* (Sydney, Australia) did not carry In992. When one copy of IS*Aba14* and the 1777-bp neighbor base sequence were lost and the other copy of IS*Aba14* was retained, a Tn*6411*-like_pWM98B_ (Tn*6411*-*aacC2*-*tmrB* region remnant-IS*Aba14*-IS*Aba14*) changed into Tn*6411*-like_pE47_ (Tn*6411*-*aacC2*-*tmrB* region remnant-IS*Aba14*) (Figure 1).

Tn*6411*-like_P8W_, Tn*6411*-like_P9W_, Tn*6411*-like_SE5430_, and Tn*6411*-like_PA34_ did not have an accessory module (the sequence of Tn*6411*-like_PA34_ was discontinuous and incomplete, and no analysis was conducted). One copy of the DR sequence (ATAT) of IS*Aba14* was found upstream of the *aacC2*-*tmrB* region remnant of Tn*6411*-like_P8W_, Tn*6411*-like_P9W_, and Tn*6411*-like_SE5430_, which enabled double copy of the IS*Aba14* sequence in the same direction to insert its Tn*6411*-like structure (Figure 1).

Compared with Tn*6411*-like_SE5430_, Tn*6411*-like_P8W_ and Tn*6411*-like_P9W_ lacked a 26-bp sequence downstream of the *aacC2*-*tmrB* region remnant, but the IRL of Tn*6411*-like_SE5430_ lacked a 14-bp sequence. TnsB binding site 1 had an 8-bp deletion as Tn*6411*_18083286_, but the difference was that TnsB binding sites 2 and 3 of Tn*6411*-like_SE5430_ were also missing. Therefore, the formation of the structure of Tn*6411*-like_P8W_ and Tn*6411*-like_P9W_ might have occurred before the formation of Tn*6411*-like_SE5430_ (Table S2).

Therefore, the *aacC2*-*tmrB* region remnant-Tn*6411* backbone was the earliest structure, and then in the evolutionary process, the element formed two evolutionary modes; one was localized in chromosome by TnsD for vertical transmission within species, and the other was localized in plasmid by TnsE for horizontal transmission between different species (Figure 2).

**Figure 2.**
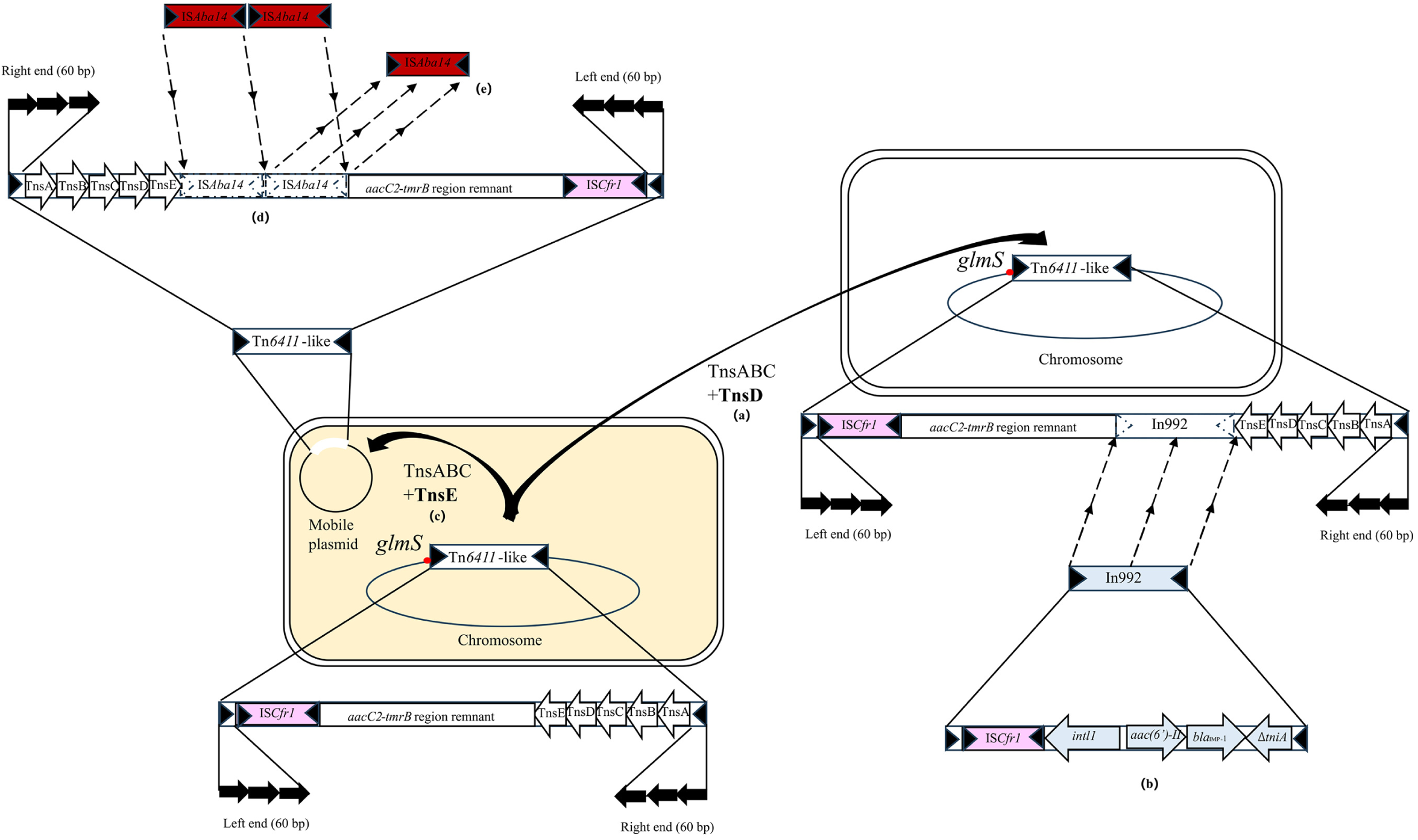
The formation process of Tn*6411* and its derived structures. The figure shows the *aacC2*-*tmrB* region remnant-Tn*6411* backbone as a local structure. IS*Cfr1* is shown in pink, attTn*7* (*glmS*) is shown as a red dot, In992 is shown in light blue, and IS*Aba14* is shown in brownish red. **(a)** A Tn*6411*-like transposon (*aacC2*-*tmrB* region remnant-Tn*6411*backbone) was localized in the chromosome by TnsD; **(b)** In992 carrying *bla*_IMP-1_ and *aac(6’’)-II* was inserted into the Tn*6411* backbone. The truncated Tn*402* transposition module in In992 had undergone the deletion of the partial TniA sequence and complete TniBQR sequence and lost its self-transfer capability, making it stable, forming Tn*6411*. **(c)** Tn*6411*-like (*aacC2*-*tmrB* region remnant-Tn*6411*) was localized in a plasmid by TnsE for horizontal transmission. **(d)** Two copies of IS*Aba14* were inserted into the Tn*6411* backbone. **(e)** One copy of IS*Aba14* and a 1777-bp neighbor base sequence were lost, and the other copy of IS*Aba14* was retained.

## Conclusion

We isolated two carbapenem-resistant *P. aeruginosa* ST865 strains with potential intrahospital dissemination risk. We found that this bacterium could capture a Tn*402*-like type 1 integron containing *bla*_IMP-1_ through the Tn*6411* transposon, which formed a stable structure, and transmit this integron vertically. It could also insert plasmids, which provide a new vector and host for the horizontal transfer of *bla*_IMP-1_. Hence, Tn*6411* and its derived structures should be closely monitored globally. At the same time, there is a need to improve disinfection methods to prevent the spread of opportunistic pathogens in hospitals.

## Author contributions

All strains were provided by China-Japan Union Hospital, Jilin University. XJG, LWZ, YLY and PC conceived, directed, and carried out the study. JYG, QLL, GJL, YS, XJ, JJ, SWS, and BWJ prepared samples for sequence analysis. JYG, LZ, and ZXW acquired samples and analyzed the data. LZ and ZXW conducted this manuscript. All authors have read and approved the final manuscript.

## Acknowledgments

We are grateful to the members of the China-Japan Union Hospital, Jilin University.

